# Sizing up phylogenetic testing in geometric morphometrics: A case study of allometry

**DOI:** 10.1101/2025.10.27.684678

**Authors:** D. Rex Mitchell, Ben Halliwell, Luke Yates, Sally Potter, Mark D. B. Eldridge, Vera Weisbecker

**Affiliations:** College of Science and Engineering, Flinders University, GPO Box 2100, Adelaide, SA 5001, Australia; Australian Research Council Centre of Excellence for Australian Biodiversity and Heritage, Wollongong, NSW 2522, Australia; School of Natural Sciences, Private Bag 55, University of Tasmania, Hobart, Tasmania, Australia; ARC Centre of Excellence for Plant Success in Nature and Agriculture, Private Bag 55, University of Tasmania, Hobart, Tasmania, Australia; School of Natural Sciences, Macquarie University, Sydney, New South Wales 2109, Australia; Australian Museum Research Institute, Sydney, New South Wales 2010, Australia

**Keywords:** Geometric Morphometrics, phylogeny, morphology, allometry, cranium, *Petrogale*

## Abstract

Accounting for phylogenetic relatedness in the analysis of shape has become a common practice, deemed necessary to factor in the non-independence between species because of common ancestry. However, when adjusting error distributions to account for relatedness, the phylogenetic-generalised-least-squares (PGLS) test can obscure an important component of variation called conservative trait correlation (CTC). This is the amount of variation in a response variable that is both attributable to a predictor variable and phylogenetically structured. If CTC represents a large amount of correlated variation, true biological associations with strong phylogenetic signal (from unrepeated evolutionary events for example) might not be supported using a PGLS. We demonstrate this effect using geometric morphometric shape analysis on 370 crania from the speciose Australian rock-wallabies (genus *Petrogale*). In this clade, well-recognised allometric patterns such as scaling of the braincase (Haller’s rule) and snout length (craniofacial evolutionary allometry) are supported using ordinary least squares (OLS) regression, but not PGLS, indicating that important between-species shape variation is lost. We then apply two methods capable of quantifying aspects of the missing variation: variation partitioning (VARPART), which estimates the proportion of variation shared between the predictor and phylogeny, and multi-response phylogenetic mixed models (MR-PMM), which identify the strength of correlation within the phylogenetic component of trait variance. Both methods show that CTC dominates the allometric shape variation in our sample, highlighting its importance in assessing phylogenetically informed models. We suggest approaches that can consider CTC become more widely used to better understand morphology and its predictors.

## Introduction

In the quantitative study of ecology and evolution, the shared ancestry of all life on Earth demands careful consideration in the choice of statistical methods that are used to examine it. This is because more closely related species tend to share more similar phenotypes than less related ones (Felsenstein 1985; Harvey & Pagel 1991). Interspecific analyses therefore carry a risk of statistical non-independence in trait variation (Garland Jr 2001; Maddison & FitzJohn 2015). To address these concerns, a statistical toolkit collectively termed “phylogenetic comparative methods” was developed (Felsenstein 1985; Grafen 1989; Garland Jr et al., 1992), and has since become a critical step of interspecific statistical analyses across much of the biological sciences.

Linear regression of comparative data has been particularly transformed by the advent of phylogenetic comparative methods, with the phylogenetic-generalised-least-squares (PGLS; Grafen 1989) method. PGLS accounts for the expected phylogenetic covariance structure among traits in a sample, by assuming a certain model of evolutionary change (such as Brownian motion) along the branches of a supplied phylogeny (Rohlf 2001; Symonds & Blomberg 2014). This is intended to reduce the relative weight of similarity of more closely related taxa to account for phylogenetic non-independence (Smaers & Rohlf 2016). The approach is now widely used across the biological sciences, and is particularly common in investigations of morphological evolution in animals. A mature and robust analytical toolkit, particularly in the R programming environment (Adams et al., 2013; Orme et al., 2013; Paradis & Schliep 2019; R Development Core Team 2023), makes it a strongly recommended analytical pathway even in the complex multidimensional contexts of geometric morphometrics (Klingenberg & Marugan-Lobon 2013; Adams 2014; Adams & Collyer 2019; Klingenberg 2022) where it is routinely applied (Rohlf 2001; Polly et al., 2016; Weisbecker et al., 2023; Mitchell et al., 2024b).

Despite the prevalence of PGLS in the analysis of morphology, an important concern is its inability to differentiate trait variation due to evolutionary chance *versus* biologically meaningful patterns. Westoby et al. (1995b) noted that a proportion of the covariation between predictor and response variables will often also correlate with phylogeny. He argued that PGLS models phylogenetic structure in only the residuals of the regression and therefore ignores structure in the predictor. Therefore, if the phylogenetic covariance structure assumed in a PGLS model overlaps with biologically meaningful covariance, such as produced by adaptation, PGLS will recognise adaptation as phylogenetic signal (resemblance of related species to each other; Blomberg & Garland Jr 2002) and thus remove an important functional component of morphological variation (Westoby et al., 1995b) (Fig. 1).

**Figure 1:**
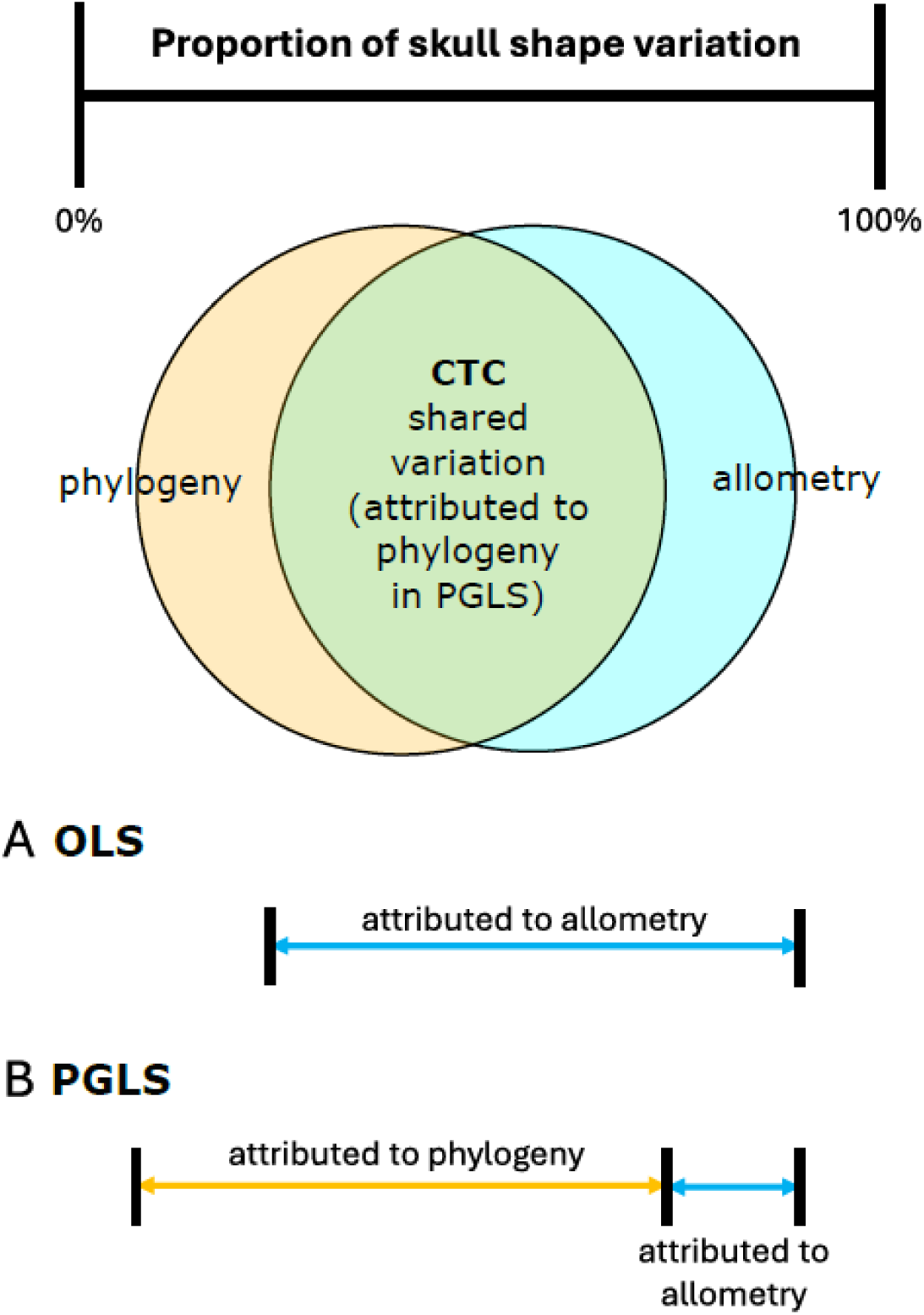
Venn diagram, modified for skull allometry from Westoby (1995b), showing the proportion of variation in skull shape correlated with both phylogeny and size (or ‘conservative trait correlation’ or CTC). The two extremes of variation attribution are presented below. (A) an OLS regression ignores relatedness, but can identify all allometric variation, (B) a PGLS accounts for relatedness, but this includes the allometric variation that is phylogenetically structured (CTC), potentially misattributing a substantial amount of allometric variation.

This inextricable quantity of variation together explained by phylogeny and variables has since come to be referred to as ‘conservative trait correlation’ (CTC) (Westoby et al., 2023; Halliwell et al., 2025). CTC is particularly relevant regarding chance evolutionary events that are often associated phylogenetic niche conservatism, because the arrival of apomorphic traits that provide increased fitness are likely to be conserved within a diversifying clade (Westoby et al., 1995b). Such chance evolutionary events can also be due to phylogenetic inertia (Felsenstein 1985) and contingency (Young et al. 2009), and these can all be highly relevant to form-function focussed hypotheses. However, because of this potential bias PGLS has with attributing variation to relatedness, phylogenetic correction can potentially miss these patterns, leading to erroneous conclusions about trait coordination and the environmental factors that govern trait evolution (Uyeda et al., 2018; Weisbecker et al., 2023).

Westoby et al.’s perspective was debated further (Harvey et al., 1995; Westoby et al., 1995a), but the concerns raised ultimately did not diminish the widespread adoption of phylogenetic comparative methods like PGLS. However, there are many cases where PGLS analyses report non-significant associations of shape with a trait, despite these associations being both expected for mechanistic reasons and significant in the non-corrected linear models (Weisbecker et al., 2021; Weisbecker et al., 2023; Mitchell et al., 2024b). A particularly clear example of a coincidence between phylogenetic structure and functional adaptation, and excellent test case, is in the allometry (size-correlated shape changes) (Gould 1966; Mosimann 1970; Klingenberg 2016) of mammalian cranial shape. For example, one of the most commonly manifested and easily recognisable patterns of allometry in the mammalian skull is that larger species, within taxa of similar ecology and behaviour, tend to have relatively smaller brains and associated braincases. This effect, known as Haller’s rule (Rensch 1948), reflects a universal pattern of negative brain size scaling with body mass and has been demonstrated among diverse animal clades (Radinsky 1981; Burger et al., 2019; Smaers et al., 2021). Other common allometric patterns include relative orbit size, and the craniofacial evolutionary allometry (CREA) pattern of facial elongation in related species with larger body sizes (Robb 1935; Emerson & Bramble 1993; Cardini & Polly 2013; Tamagnini et al., 2017; Mitchell et al., 2024b). However, in a PGLS context, these expected patterns are not always retrieved even when they appear both in visual inspection and in a non-corrected model because animal body size is also often phylogenetically structured.

Cope’s rule, for example, is the pattern whereby body size tends to be larger in more recently evolved species (Cope 1896; Hone & Benton 2005). There are also many cases where some branches of a clade will solely contain all of the largest species represented, or solely all of the smallest species (Mitchell et al., 2024b). These represent the kinds of instances of unreplicated, singular events to which phylogenetic testing can be vulnerable (Uyeda et al., 2018).

As would be expected from a misattribution of CTC as a purely phylogenetic phenomenon, support for Haller’s rule and other widely-known allometric patterns frequently vanishes upon the inclusion of phylogeny into models (Mitchell et al., 2024b). Because autocorrelation between size and relatedness is so common in biology, the confounding effects of phylogeny in PGLS allometry models is likely a widespread occurrence. This was shown in similar tests focused on a genus of Australian rock-wallabies (Mitchell et al., 2024a), which led to the suggestion of ignoring phylogenetic structuring from tests of evolutionary allometry (see similar rationale in the methods of Weisbecker et al., 2023). But in reality, neither OLS or PGLS are correct approaches for analysing phenotypic variation where phylogenetic signal and true adaptation might be conflated. OLS ignores the influence of relatedness on error distributions, while PGLS can either diminish, or mask altogether, significant correlations between predictor and response variables that are phylogenetically structured. A modified version of Westoby et al. (1995b)’s Venn diagram neatly encapsulates the problem (Fig. 1). It is here presented in the context of skull allometry, but theoretically holds true for any quantitative variables that have a significant phylogenetic signal. The diagram highlights the information potentially lost when applying PGLS.

There are two established methods that are excellent candidates for distinguishing the relationship between variables and relatedness in comparative morphological studies. The first is variation partitioning (VARPART) (Legendre et al., 2012; Legendre & Legendre 2012). This method has been used previously in geometric morphometrics to assess the joint contributions of predictor variables on shape (Mitchell et al., 2020; Viacava et al., 2020; Pierre et al., 2025), and to partition shape variation correlated with phylogeny and allometry (Piras et al., 2010). The second method is the multi-response phylogenetic mixed effects (MR-PMM) modelling (Westoby et al., 2023; Halliwell et al., 2025), which allows investigation of the relationships between predictor and response variables within both phylogenetic and non-phylogenetic components of variance.

Here, we used landmark data from the crania of a speciose and geographically widespread Australian genus *Petrogale* (Macropodidae) rock-wallabies (Mitchell et al., 2024a) as an ideal model system to assess the use of VARPART and MR-PMM in clarifying the shared phylogenetic component of allometry. After a recent and rapid diversification (estimated 0.5-7mya; Potter et al., 2012), *Petrogale* rock-wallabies today comprise 17 species ranging across an order of magnitude in body mass, from ∼1.5 kg to ∼12 kg (Maynes 1989; Churchill 1997; Johnson & Delean 1999; Telfer & Bowman 2006; Richardson 2012) including two dwarf species and a species complex of five sub-species. Having been used as a model system in genetic research, the phylogeny of this genus is well-studied (Potter et al., 2012; Potter et al., 2017; Potter et al., 2022). The previous research on cranial shape of this genus (Gomes Rodrigues et al., 2016; Mitchell et al., 2024a) shows that rock-wallabies have clearly discernible allometric patterns predicted by Haller’s rule (relative braincase size) and CREA (relative facial elongation) (Cardini et al., 2015; Tamagnini et al., 2017) (Fig. 2A).

**Figure 2:**
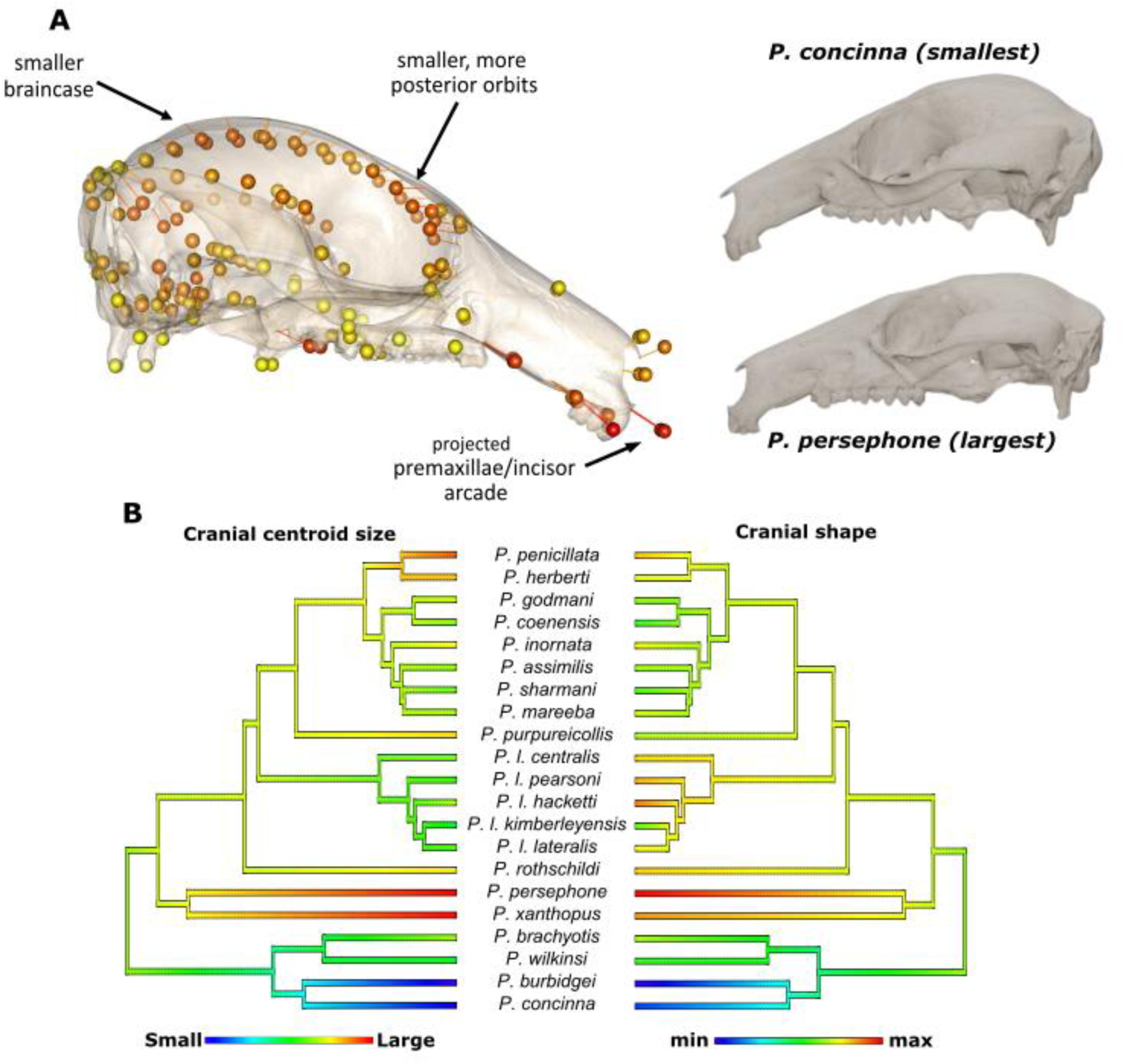
(A) Allometry in *Petrogale* crania (Mitchell et al., 2024a). Larger species demonstrate the labelled features. The mesh represents shape predicted for smallest cranial sizes, while the points represent predicted shape for largest sizes (B) Cranial size and cranial shape are both phylogenetically structured in the *Petrogale* genus, with close correspondence. Shape represented by the shape regression score of a regression with centroid size.

Both skull size and shape are also phylogenetically structured and highly congruent in their phylogenetic distributions (Fig. 2B). Two monophyletic clades representing the smallest and largest extremes of size can be viewed as unreplicated singular events which might confound PGLS models of allometry as variation attributable to CTC. We expected our two methods to identify the CTC as a shared proportion of shape variation that can be explained by both relatedness and size.

## Material and methods

The dataset of *Petrogale* crania (Mitchell et al., 2024a) comprised 370 individuals representing all 17 recognised species. The sample of 83 *P. lateralis* specimens were also partitioned into their five recognised subspecies, because of their high chromosomal variation (Potter et al., 2017).

The processed landmark data included 102 fixed and 48 semi-sliding landmarks distributed across the whole crania (Supplementary Table S1) were imported into R (v. 4.4.2) (R Development Core Team 2023) via R Studio (v.2025.05.0) and analysed using the “geomorph” package (Adams et al., 2013). An initial Procrustes superimposition was then performed on the raw coordinates, in order to remove degrees of freedom and variation attributable to translation, rotation, and size (Rohlf & Slice 1990). We additionally removed variation due to asymmetry using the ‘bilat.symmetry’ function. The resulting Procrustes coordinates were used in the subsequent analyses.

To estimate the phylogenetic tree, we used a single individual from each species, as well as a single individual representing each sub-species of *P. lateralis*, and *Dendrolagus lumholtzi* as the outgroup (n=22) (Supplementary Table S2). We used the phylogenetic tree topology from previous research (Potter et al., 2017; Potter et al., 2022) as input and estimated divergence times using MCMCTree (Rannala & Yang 2007) using the concatenated dataset of 1961 nuclear loci (Potter et al., 2017). Soft constraints were used to estimate divergence times based on previous fossil calibrated phylogenetic results (Potter et al., 2012). We set the root age at 6-11.3 million years ago (mya) and included four secondary calibrations: one for the *brachyotis* group node (1.9-5.6 mya); one for the penicillata group excluding *P. herberti* and *P. penicillata* (900,000-2.7 mya); one for the clade including *P. rothschildi*, *P. lateralis*, *P. purpureicollis* and the *penicillata* group (4.4-5.5 mya); and one including all species except those in the *brachyotis* group (1.9-8.7 mya). MCMCtree was run using the HKY nucleotide substitution model, 0.5 alpha for gamma rates at sites and four discrete gamma categories. We used an independent rates model, the approximate likelihood calculation (dos Reis & Yang 2011) and included missing sites in the analysis. The analysis was run sampling every 1000 generations for a total of 10,000 samples after a burnin of 1 million generations. We checked the analysis had reached convergence using Tracer v1.7.2 (Rambaut et al., 2018) with effective sample sizes >200 indicative of stationarity and a suitable run length.

### Standard geometric morphometric analyses

To represent cranial size in tests of allometry, we used cranial centroid size (the square root of the sum of squared distances of all the landmarks of an object from their centre of gravity). A test for phylogenetic signal is often conducted as a means of justifying phylogenetically informed analysis (Symonds & Blomberg 2014). We therefore first estimated the phylogenetic signal for mean taxon cranial shape and mean taxon cranial size using the ‘physignal’ function. We then computed an initial Procrustes regression of shape ∼ log(size) on the whole sample including within-species replicates, using the ‘procD.lm’ function with 1000 iterations, and then performed the same test using only the mean values for the 21 taxa. Following this, we applied a phylogenetic-generalised-least-squares (‘procD.pgls’ function; 1000 iterations), again using the mean shape and size data from each taxon.

### Variation Partitioning (VARPART)

In order to evaluate the degree of shared influence of cranial allometry and phylogenetic relatedness on cranial shape, we performed a variation partitioning analysis (VARPART: Legendre et al., 2012; Legendre & Legendre 2012), using the “vegan” package (Oksanen et al., 2013). We did this on the entire sample, in order to include intraspecific variation in our models (Garamszegi & Moller 2010). To include phylogenetic structure as a variable in VARPART, we used the ‘pcoa’ function from the “ape” package to extract phylogenetic eigenvectors from a pairwise phylogenetic distance matrix generated from the tree (Diniz-Filho et al., 1998; Piras et al., 2010; Guénard et al., 2013; Diniz-Filho et al., 2014; Shapiro et al., 2025). Cranial size and the full set of phylogenetic eigenvectors were modelled as predictors of cranial shape using the ‘varpart’ function. The output gives the adjusted R-squared values for all fractions of a Venn diagram, for which we obtained all testable R-squared values and p-values using a redundancy analysis with the ‘rda’ function. This approach provided an estimate of how much variation in the sample was explained by both size and relatedness. We focussed on adjusted R-squared values because the standard R-squared value computed in RDA is a biased estimator with larger values often found alongside the inclusion of more explanatory variables in the model (Peres-Neto et al., 2006).

### Multi-response phylogenetic mixed models (MR-PMMs)

We used Bayesian MR-PMM to decompose covariances between cranial size and shape into phylogenetic and non-phylogenetic components (Halliwell et al., 2025). To represent size-related shape variation (see Discussion), we used the Shape Regression Score (SRS), obtained by projecting the shape data onto a normalised regression vector for the size variable. The SRS is single measure of shape changes predicted by the regression model of shape ∼ log(size), and includes the residual variation (Drake & Klingenberg 2008). We elected to use the SRS instead of a different ordination, such as the first principal component of shape, because allometric variation can often be distributed across several principal components (Mitchell et al., 2024a), while the SRS factors in all of the shape-correlated variation. Specifically, we fit both traits (size and SRS) jointly as response variables, using correlated random effects to specify phylogenetic and non-phylogenetic between-species covariance matrices. We modelled multi-level structure in the data by fitting random intercepts across species (accounting for within-species replication). Retaining observation-level data also allowed us to estimate within-species residual (co)variances, and more accurately estimate phylogenetic signal (Cinar et al., 2021). All MR-PMM were fit using the brms package (Bürkner 2017) in R (also version 4.4.2).

Patterns of cranial size and shape evolution across the phylogeny were visualised (Figure 2) using character mapping in the phytools package (Revell 2012).

## Results

### Standard Geometric Morphometrics tests

We identified significant phylogenetic signal for mean taxon values of cranial shape (K=0.657, p=0.001) and cranial centroid size (K=1.126, p=0.002), which would typically encourage the use of phylogenetically-informed tests. Regression of shape and cranial centroid size revealed significant evidence of allometry (Table 1). Significant effects of allometry remained when testing taxon means without the phylogeny (OLS); however, when phylogeny was included in a phylogenetic regression, allometry was no longer significant, consistent with results previously obtained for this data in Mitchell et al. (2024a).

**Table 1:** Results for each of the tests frequently used to test the effect of allometry in Geometric Morphometrics research.

| Test | df | Rsqr | F | p |
| --- | --- | --- | --- | --- |
| **(cranial shape ~ log(cranial size))** |  |  |  |  |
| OLS full sample | 1,369 | 0.173 | 76.851 | **0.001** |
| OLS means | 1,19 | 0.370 | 11.169 | **0.001** |
| PGLS means | 1,19 | 0.057 | 1.149 | 0.279 |

### Variation Partitioning (VARPART) analysis

VARPART explained an estimated 47.4% of total shape variation. The analysis identified significant influences of both cranial size and phylogenetic relatedness on cranial shape, across both pure and total fractions (Table 2; Fig. 3). An estimated 10% of total shape variation was shared between cranial size and relatedness, while a smaller proportion of 7.0% was identified as purely allometric.

**Figure 3:**
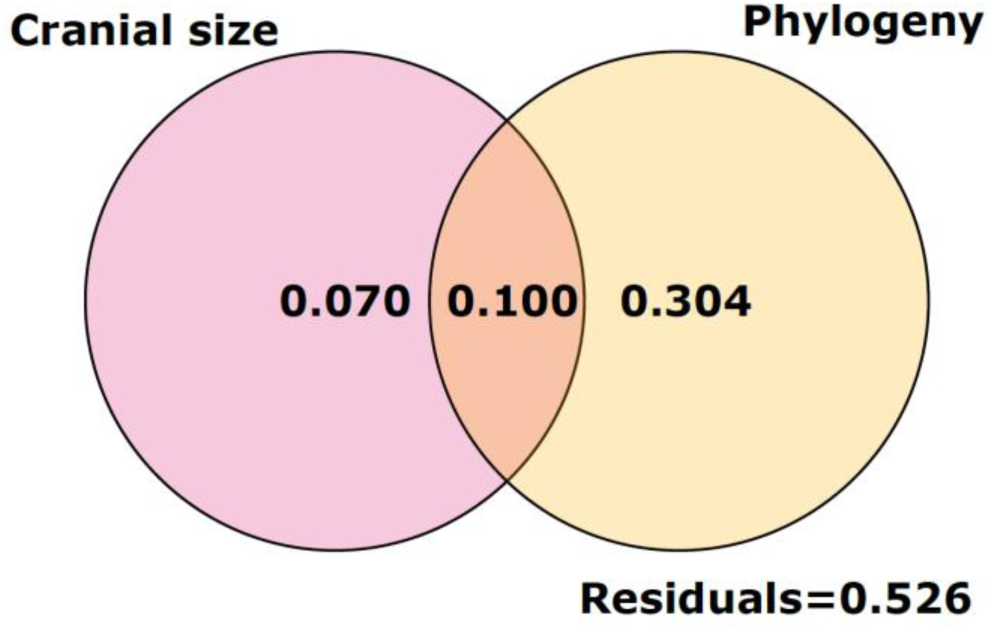
VARPART Venn diagram, indicating an estimated 10% of shape variation is shared between Phylogeny and allometry.

**Table 2:** Variation partitioning analysis (VARPART) results of the effects of allometry on cranial shape. Bold values indicate significance (*P* < 0.05).

| **Allometry** | **df** | **R<sup>2</sup>** | **Rsqr.adj** | **P** |
| --- | --- | --- | --- | --- |
| Full model | 21 | 0.505 | 0.474 | **0.001** |
| Total fractions |  |  |  |  |
| Cranial size | 1 | 0.173 | 0.171 | **0.001** |
| Phylogeny | 20 | 0.436 | 0.404 | **0.001** |
| Pure fractions |  |  |  |  |
| Cranial size | 1 | 0.068 | 0.070 | **0.001** |
| Phylogeny | 20 | 0.331 | 0.304 | **0.001** |
| Estimated shared | 0 |  | 0.100 | Untestable |
| Residuals |  |  | 0.526 |  |

### Multi-response phylogenetic mixed models

The phylogenetic mixed-effects model fit to the whole sample revealed strong phylogenetic signal for both body size and shape across species (Table 3). The phylogenetic between-species correlation (i.e., CTC) between size and shape was strongly positive (Table

**Table 3:**
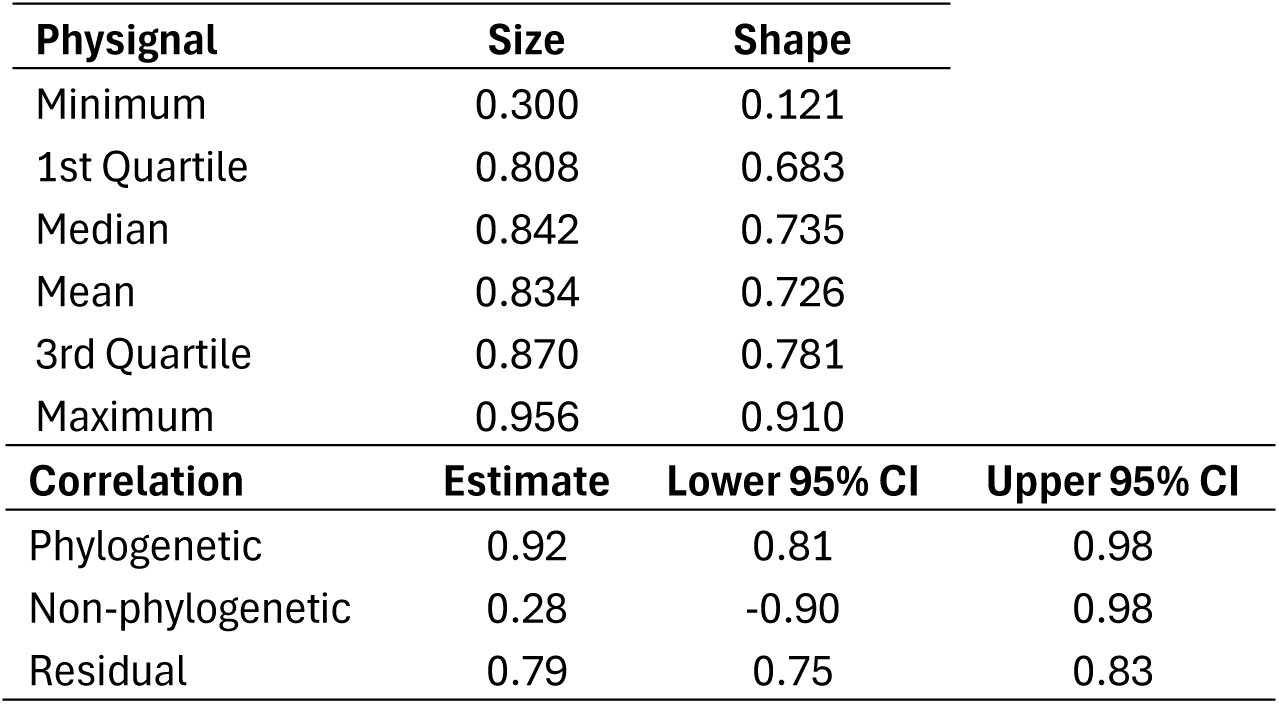
Physignal ranges > 0 indicate significant phylogenetic signals for size and shape. Correlation estimates and confidence intervals >0 show significant positive correlations of size and shape along PC1 for Phylogenetically-informed interspecific (Phylogenetic) and intraspecific (Residual) allometry.

3; Fig. 4A), indicating that larger-bodied species tend to exhibit allometric shifts towards the maximum of the shape regression score across the phylogeny. While a positive non-phylogenetic between-species relationship was also inferred (Fig. 4A), a very wide credible interval suggests limited evidence for phylogenetically independent allometric variation between species. There was, however, a strong residual allometric effect within species, such that individuals of a species with relatively larger skulls also displayed relatively high SRS values (Table 3). A variance decomposition (Fig. 4B) shows that both size and SRS of shape have a high proportion of variance explained by phylogeny, with only a very small amount being non-phylogenetic. The SRS has a greater proportion of residual variance than cranial size.

**Figure 4:**
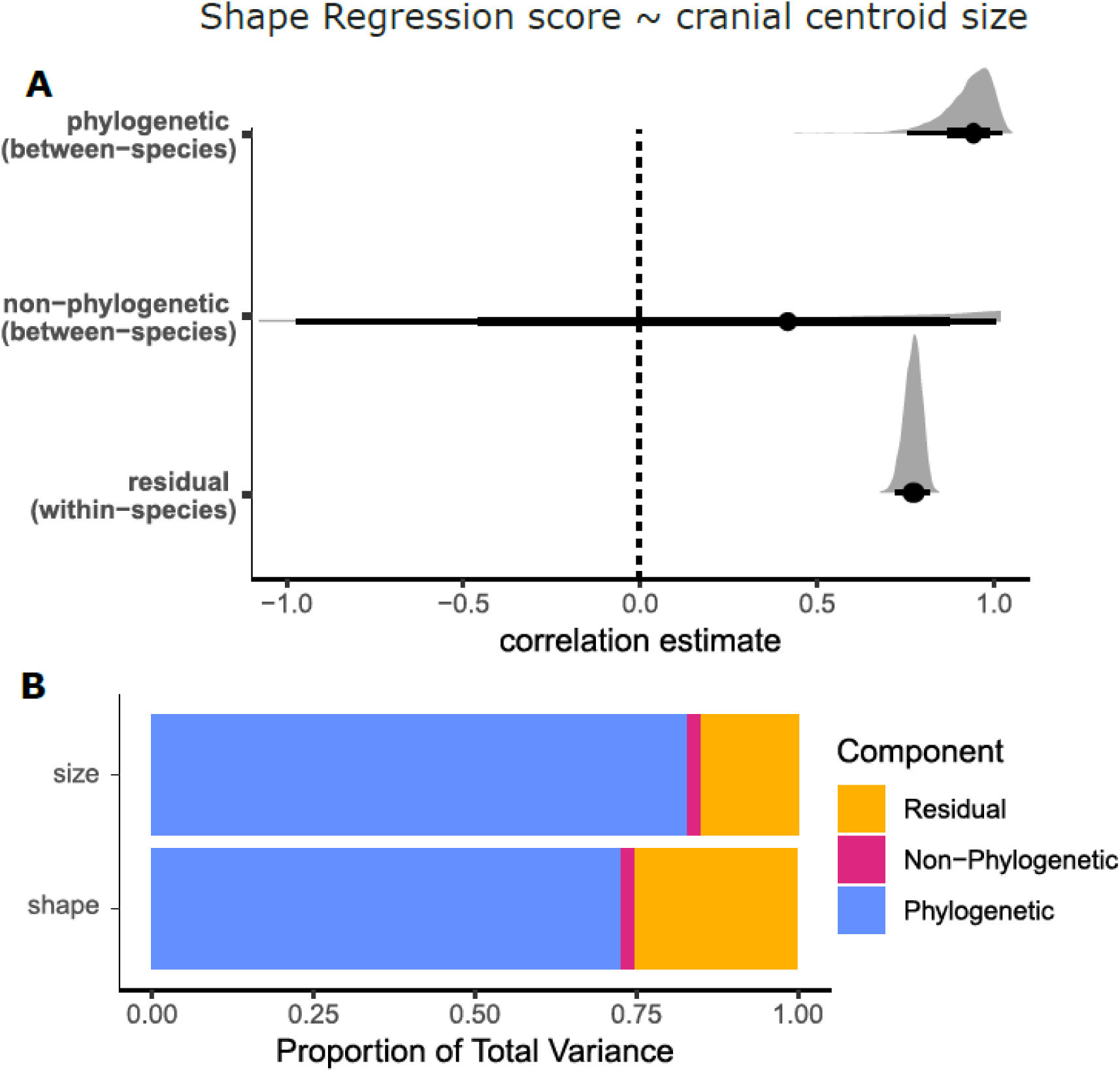
(A) Venn diagram of VARPART analysis indicates allometric variation both correlated with phylogeny and independent of phylogeny. (B) The MR-PMM shows a significant positive correlation between cranial shape and cranial centroid size within a phylogenetic context (confidence interval well above zero), while there was no evidence for such a correlation outside of the context. There is also a significant residual correlation indicating within-species allometry.

## Discussion

In the study of morphology, size-related constraints on anatomy, physiology, behaviour, and ecology mean that allometric variation is almost certain to exist to some degree in any study sample that contains appreciable size variation. Allometric variation therefore serves as a useful indicator for testing the capability of different approaches to detect known correlations within a phylogenetic context. In our test case of *Petrogale* cranial variation, we showed that obvious differences in cranial shape according to size were reflected as significant allometric patterns in an OLS analysis. The shape differences of the genus are dominated by relative brain size and face length (Fig. 2A), consistent with predictions of Haller’s rule and CREA (Rensch 1948; Cardini & Polly 2013; Mitchell et al., 2024b). However, no significant evidence of these well-known patterns was produced using a PGLS. We expect this to be due to the monophyletic branches within which almost all cases of extreme values of a cranial size and associated shapes are represented (Fig. 2B). This allometric variation contributed by unreplicated, singular events would be effectively excluded from estimates of allometric relationships, due to the adjustment of error distributions expected from phylogenetically structuring within the PGLS framework. The PGLS was not, as would typically be interpreted, identifying no influence of allometry across the clade, but merely did not retrieve significant associations in the phylogenetically unstructured data (i.e., the pure blue fraction of Fig. 1). This reinforces existing concerns that PGLS testing alone is likely inadequate for assessing allometry in most cases (Westoby et al., 1995b; Mitchell et al., 2024a; Mitchell et al., 2024b), and we suggest more detailed approaches to account for the missing portion of variation explained by both allometry and phylogeny. These arguments and their rationale are also applicable to the analysis of any quantitative species traits that display strong phylogenetic signals.

The significant phylogenetic signal in both response and predictor variables means that OLS regression is not appropriate, because the errors are not independent due to relatedness (Klingenberg & Marugan-Lobon 2013). However, a PGLS is also clearly inappropriate, because the process of accounting for relatedness in the error structure, while failing to model phylogenetic signal in the predictor, can mask the significance of known patterns of allometry. Both VARPART and MR-PMM appear to address this issue and yield overall similar results by acknowledging the component of variation shared by both the predictor and phylogenetic structure. However, there are key differences in the quantities these methods estimate, affecting model inference, and some important considerations for both approaches.

In VARPART, variance components are derived as differences in the outputs of a sequence of model fits rather than being estimated directly as model parameters (Peres-Neto et al., 2006; Legendre et al., 2012). This approach is computationally efficient, scaling easily to large multivariate data sets and generating the same decomposition structure irrespective of the number of response variables. However, without a statistical model for the variance components themselves, statistical inferences are limited to permutation tests for the individual predictor sets. The inclusion of phylogenetic eigenvectors as fixed predictors in the underlying regressions avoids the complexity of modelling correlated residuals (as implemented in MR-PMM), but decouples the vectors from their associated eigenvalues weakening their relationship to the assumed evolutionary model. This inability to properly account for and decompose variance across hierarchical data structures is a key limitation of VARPART approach.

Despite this, VARPART was able to identify significant associations of both size and phylogeny on cranial shape and provide an estimation of the proportion of total shape variation shared by both variables. This conceptually addresses the problem originally posed by Westoby (1995b) in providing an approximation for the missing shared value (Fig. 1). In the case of our sample, we found ∼10% of total shape variation was shared between cranial size and relatedness, representing more than half of all allometry detected in the sample. This can be a valuable metric to assess phylogenetic niche conservatism or morphological contingencies; however, VARPART is unable to indicate whether this shared variation bears a significant impact. This is because a limitation of VARPART is that the significance of estimated shared proportions of variation are untestable (Cubo et al., 2008; Osburn et al., 2023). Additionally, VARPART was able to estimate the proportion of allometric shape variation that is independent of phylogeny and report a measure of significance for it.

However, the method has no means of discerning whether this is intraspecific allometry (ontogenetic/static) or represents some additional interspecific patterns that are independent of phylogenetic structure.

The MR-PMM is a more powerful approach that provides more detailed outputs of parameter estimates and their uncertainty. It represents a unified statistical model which explicitly parameterises the variances and covariances between all traits (ordinated shape components and body size). The model directly estimates separate (co)variance components for each specified source, here phylogenetic and non-phylogenetic variation at the species level, plus within-species variation (Westoby et al., 2023; Halliwell et al., 2025). This direct parameterization enables statistical inference on the variance components themselves, including credible intervals and hypothesis tests about the relative importance of evolutionary versus other processes. While VARPART aggregates variance decomposition across all response variables, MR-PMM decomposes covariances for individual ordinated components, which may be advantageous when components have distinct biological interpretations and differing phylogenetic signals.

Similar to VARPART, the MR-PMM identified a significant correlation between cranial size and shape, but in this case, it identified the relationship within a phylogenetic context. In other words, the MR-PMM can confirm the significance of a shared fraction of allometry and phylogeny that was also estimated – but untestable – using VARPART. The MR-PMM also differed from VARPART in being able to identify the significant role of intraspecific (i.e., ontogenetic/static) allometry in the sample, using hierarchical random effects. A variance decomposition was further able to demonstrate the dominance of phylogenetic relatedness in the sample. However, the finer resolution provided by MR-PMM comes at the cost of computational complexity, creating challenges for both fitting and interpretation that may require focusing on the leading components. This is because there is a risk of overparameterisation when modelling high-dimensional multivariate data. Many of the questions that commonly arise in geometric morphometrics involve multivariate data (Monteiro 2013; Adams & Collyer 2018), but while VARPART is able to include all shape data into the model, the shape data in MR-PMM relies on a decomposition down to a single variable, or at least few variables. There is nothing wrong in principle with including multiple, shape-correlated PC components as response variables in MR-PMM where clear interpretation allows. However, there might be an issue for some data sets where size-correlated variation related to the predictor variable is spread more evenly across many principal components (Weisbecker et al., 2019). Because the *Petrogale* dataset is known to have allometry distributed across at least two principal components (Mitchell et al., 2024a), we chose to decompose the multivariate shape data to a single measure, in the form of allometry-correlated shape regression scores. These important considerations are also relevant to other multivariate data, such as climate or spatial data.

For the shared fraction of variation, both allometry and phylogeny are equally feasible – and perhaps inextricable – hypotheses to explain the shape patterns it describes (Mitchell et al., 2024a). For these proportions of variation, researchers must therefore rely on an objective appraisal of the knowledge of other biological systems and well-supported patterns to interpret the reasons behind singular events. For example, the monophyletic clade of the two dwarf species of rock-wallabies might indicate historical niche partitioning (Sanson et al., 1985), Bergmann’s rule (Bergmann 1847; McNab 1971; Meiri 2010; Clauss et al., 2013), or other unknown contingencies. Yet, regardless of the causes behind their small sizes, the two smallest species exhibit cranial shape expected from Haller’s rule and also the CREA pattern of relatively shorter faces in smaller species within certain clades (Cardini & Polly 2013; Cardini et al., 2015). These species do not represent morphological aberrations to expected shape. Allometry-related shape variation is almost certainly the major driver of shape differences in the phylogeny-shared proportion of variation across the genus, previously described in detail as a combination of Haller’s rule and allometry of functional demands relating to bite force optimisation (Mitchell et al., 2024a). A result of no significant allometry from a PGLS would in all likelihood be an inaccurate finding to report, which emphasises the need to acknowledge CTC in the study of morphology.

## Conclusion

When a significant phylogenetic signal is found in the variables of interest, neither OLS or PGLS are appropriate approaches to analyse morphological variation, because OLS does not consider non-independence of species and PGLS biases results by masking conservative trait correlation. VARPART provides a means of using multivariate data to identify significant roles of predictors and phylogeny, and can also provide a simple estimate the CTC. However, it is limited in its ability to decompose variance across hierarchical data, and to discern the significance of the CTC relationship. MR-PMM provide a more powerful, quantitatively rigorous means of discerning this relationship and can also identify the role of intraspecific variation, but it can be limited by the need to ordinate complex multivariate data. With careful treatment of these considerations, these two methods can address common problems encountered with the PGLS, through identifying significant relationships that are clearly present in the sample, whether independent of phylogenetic structuring or as part of CTC. This approach offers a more detailed view of evolutionary history that accounts for clade-specific traits. These findings are also highly relevant and applicable to the analysis of other quantitative variables tested in the study of morphology.

## Acknowledgements

We would like to thank Gavin Dally and Barry Russell of the MAGNT, Kenny Travouillon of the WAM, David Stemmer of the SAM, Sandy Ingleby of the AM, Heather Janetzki of the QM, and Leo Joseph of the ANWC (https://ror.org/059mabc80) for access to their specimens; and Erin Mein for scanning several specimens from the MAGNT. Rex Mitchell and Vera Weisbecker were funded by the Australian Research Council Centre of Excellence for Australian Biodiversity and Heritage CE170100015 and Vera Weisbecker was also funded by ARC Future Fellowship FT180100634.

## Data Availability

Data and code available at https://github.com/DRexMitchell/Petrogale_phylotests

