## Supplementary Table S2 for "Sizing up phylogenetic testing in geometric morphometrics: A case study of allometry"

**Supplementary Table 1** Taxon information for each individual, including: sample ID, species/sub-species/race, locality, latitude and longitude and sample ID associated with short read archive project PRJNA360868 where raw reads are available.

| **Sample ID** | **Tissue ID** | **Museum Voucher** | **Species** | **Locality** | **State** | **Latitude** | **Longitude** | **SRA Sample ID** |
| --- | --- | --- | --- | --- | --- | --- | --- | --- |
| S612 |  | CM13533 | *assimilis* | Hillsborough | Queensland | -20.10 | 147.04 | [SAMN06233289](https://www.ncbi.nlm.nih.gov/biosample/SAMN06233289) |
| S296 |  | CM15326 | *brachyotis brachyotis* | Beverley Springs | Western Australia | -16.51 | 125.33 | [SAMN06233290](https://www.ncbi.nlm.nih.gov/biosample/SAMN06233290) |
| S280 |  | CM15326 | *burbidgei N* | Crystal Cr., Mitchell Plateau | Western Australia | -14.50 | 125.79 | [SAMN06233295](https://www.ncbi.nlm.nih.gov/biosample/SAMN06233295) |
| S865 |  | CM13479 | *coenensis* | Musgrave | Queensland | -14.80 | 143.43 | [SAMN06233297](https://www.ncbi.nlm.nih.gov/biosample/SAMN06233297) |
| S315 | EBU35578 | none | *concinna canescens* | Mt Borrodaile | Northern Territory | -12.05 | 132.90 | [SAMN06233298](https://www.ncbi.nlm.nih.gov/biosample/SAMN06233298) |
| S657 |  | CM13466 | *godmani* | Bathurst Heads | Queensland | -14.31 | 144.21 | [SAMN06233301](https://www.ncbi.nlm.nih.gov/biosample/SAMN06233301) |
| S737 |  | CM13506 | *herberti* | Mt Donneybrook | Queensland | -22.51 | 146.74 | [SAMN06233304](https://www.ncbi.nlm.nih.gov/biosample/SAMN06233304) |
| S136 |  | CM15211 | *inornata* | Maiden Mountain | Queensland | -19.93 | 147.87 | [SAMN06233306](https://www.ncbi.nlm.nih.gov/biosample/SAMN06233306) |
| S217 |  | CM15310 | *lateralis hacketti* | Westall Island, Recherche Archipelago | Western Australia | -34.08 | 122.97 | [SAMN06233308](https://www.ncbi.nlm.nih.gov/biosample/SAMN06233308) |
| S970 |  | CM16823 | *lateralis lateralis* | Nangeen Hill | Western Australia | -31.83 | 117.68 | [SAMN06233309](https://www.ncbi.nlm.nih.gov/biosample/SAMN06233309) |
| S955 |  | CM24567 | *lateralis centralis* | Telegraph Station, Alice Springs | Northern Territory | -23.67 | 133.88 | [SAMN06233312](https://www.ncbi.nlm.nih.gov/biosample/SAMN06233312) |
| S997 |  | CM24572 | *lateralis pearsoni* | Wedge Island | South Australia | -35.16 | 136.46 | [SAMN06233313](https://www.ncbi.nlm.nih.gov/biosample/SAMN06233313) |
| S1256 | EBU47376 | none | *lateralis kimberleyensis* | Erskine Range, West Kimberley | Western Australia | -17.81 | 124.33 | [SAMN06233315](https://www.ncbi.nlm.nih.gov/biosample/SAMN06233315) |
| S436 |  | CM10492 | *mareeba* | Anthill Creek | Queensland | -18.39 | 145.14 | [SAMN06233318](https://www.ncbi.nlm.nih.gov/biosample/SAMN06233318) |
| S766 | EBU35579 | CM16835 | *penicillata* | Jenolan Caves | New South Wales | -33.82 | 150.03 | [SAMN06233319](https://www.ncbi.nlm.nih.gov/biosample/SAMN06233319) |
| S869 | EBU47379 | AM37280 | *persephone* | Gloucester Island | Queensland | -20.04 | 148.45 | [SAMN06233321](https://www.ncbi.nlm.nih.gov/biosample/SAMN06233321) |
| S888 | EBU47380 | CM16832 | *purpureicollis* | Dajarra | Queensland | -21.67 | 139.29 | [SAMN06233323](https://www.ncbi.nlm.nih.gov/biosample/SAMN06233323) |
| S204 | EBU47381 | CM15324 | *rothschildi* | Rosemary Island, Dampier Archipelago | Western Australia | -20.49 | 116.59 | [SAMN06233325](https://www.ncbi.nlm.nih.gov/biosample/SAMN06233325) |
| S110 |  | CM15203 | *sharmani* | Mt. Claro | Queensland | -18.87 | 145.73 | [SAMN06233328](https://www.ncbi.nlm.nih.gov/biosample/SAMN06233328) |
| S267 |  | CM15172 | *wilkinsi* | Butterfly Gorge, Douglas R. | Northern Territory | -13.75 | 131.58 | [SAMN06233329](https://www.ncbi.nlm.nih.gov/biosample/SAMN06233329) |
| S384 | RW307 | CM05793 | *xanthopus xanthopus* | Cootawundi Station | South Australia | -31.05 | -142.08 | [SAMN06233334](https://www.ncbi.nlm.nih.gov/biosample/SAMN06233334) |
| S1475 |  | none | *Dendrolagus lumholtzi* | Old Palmerston Hwy, Atherton Tablelands | Queensland | -17.56 | 145.61 | [SAMN06233335](https://www.ncbi.nlm.nih.gov/biosample/SAMN06233335) |
